# Unreviewed science in the news: The evolution of preprint media coverage from 2014-2021

**DOI:** 10.1101/2023.07.10.548392

**Authors:** Alice Fleerackers, Kenneth Shores, Natascha Chtena, Juan Pablo Alperin

## Abstract

It has been argued that preprint coverage during the COVID-19 pandemic constituted a paradigm shift in journalism norms and practices. This study examines whether, in what ways, and to what extent this is the case using a sample of 11,538 preprints posted on four preprint servers—bioRxiv, medRxiv, arXiv, and SSRN—that received coverage in 94 English-language media outlets between 2014–2021. We compared mentions of these preprints with mentions of a comparison sample of 397,446 peer reviewed research articles indexed in the Web of Science to identify changes in the share of media coverage that mentioned preprints before and during the pandemic. We found that preprint media coverage increased at a slow but steady rate pre-pandemic, then spiked dramatically. This increase applied only to COVID-19-related preprints, with minimal or no change in coverage of preprints on other topics. In addition, the rise in preprint coverage was most pronounced among health and medicine-focused media outlets, which barely covered preprints before the pandemic but mentioned more COVID-19 preprints than outlets focused on any other topic. These results suggest that the growth in coverage of preprints seen during the pandemic period may imply a shift in journalistic norms, including a changing outlook on reporting preliminary, unvetted research.

## 1.0 Introduction

On January 10, 2020, the World Health Organization published its first set of guidelines for preventing and controlling a suspected “novel coronavirus (nCoV)” (WHO, 2020). Soon journalists found themselves plunged into an unexpected crisis, with an out-of-control, little understood infectious disease, and an influx of new scientific information to sift through and report on. Without much peer reviewed literature to go on—especially in the early stages of the pandemic—many turned to preprint servers to share urgent new information with the public (Fraser et al., 2021). The ensuing media coverage of preprints seen during the pandemic has since been described as a complete rupture from past reporting practices (e.g., Burke, 2021; Makri, 2021). Yet, empirical evidence supporting this assertion is lacking. As noted in previous research, there is currently an absence of longitudinal investigations that examine preprint coverage over time and which assess the impact of COVID-19 on journalistic practices and norms (Fleerackers et al., 2023). This study fills this gap by examining how media coverage of preprints has evolved, both qualitatively and quantitatively, in the lead up to, and during the first year of, the COVID-19 pandemic. Using Altmetric data, it examines changes in the volume and nature of media coverage of 11,538 preprints posted between 2013 and 2021 on bioRxiv, medRxiv, arXiv, and SSRN— four of the most actively used servers used to share COVID-19-related research (Waltman et al., 2021).

## 2.0 Literature review

### 2.1 Preprint media coverage before and during the COVID-19 pandemic

Preprints have been used extensively in physics, math, and computational science since arXiv launched in 1991. However, scientists in the biological and medical fields have been more reluctant to do so—that is, until recently (Puebla et al., 2021). The early months of the pandemic saw a sharp increase in the volume of available COVID-19-related preprints (Funk, 2023; Horbach, 2020), with preprint servers such as medRxiv and bioRxiv becoming key disseminators of pandemic research (Else, 2020; Vergoulis et al., 2021). One study (Kousha & Thelwall, 2020) found that preprints posted to arXiv bioRxiv, medRxiv, and SSRN comprised 13.26% of the COVID-19 literature during March–April 2020, while an analysis by Fraser et al. (2021) found that preprints posted to 16 servers (including the four examined in this study) comprised almost 25% of the COVID-19-related research available from January–October 2020.

COVID-19-related preprints also gained traction within news media, receiving coverage in diverse media outlets around the world (Fleerackers et al., 2021; Massarani et al., 2021; Massarani & Neves, 2021; Simons & Schniedermann, 2023; van Schalkwyk & Dudek, 2022). One study found that more than a quarter of COVID-19-related bioRxiv and medRxiv preprints were mentioned in at least one media story during the pandemic, while only about 1% of those on other topics received media coverage (Fraser et al., 2021). Some journalists reported adopting novel practices to report on these unreviewed studies, something they said they had never done before (Fleerackers et al., 2022a; Massarani et al., 2021).

This media coverage of preprints seen during the COVID-19 pandemic has been described by some journalists as a “paradigm shift” (Fleerackers et al., 2022a). Yet, while studies conducted during the COVID-19 pandemic provide important evidence into how journalists covered preprints during the evolving health crisis, little is known about whether journalists have covered preprints on other topics or during other communication contexts. For example, Fraser et al. (2021)’s widely cited study is often described as providing evidence that “During the pandemic, journalists…paid increased attention to preprints” (Kwon, 2021), but the authors did not compare pandemic preprint coverage to pre-pandemic levels. Instead, they provided evidence that COVID-19-related preprints received an outsized amount of media attention, *relative to those on other topics posted to bioRxiv and medRxiv during the same time period—*but not relative to preprints posted during different time periods or on different servers (Fraser et al., 2021). One recent study begins to fill this gap through an examination of coverage of preprints by seven German newspapers from 2018-2021 (Simons & Schniedermann, 2023). The authors identified low and stable rates of coverage leading up to the pandemic, followed by a major surge in 2020 and 2021 that was driven by COVID-19-related preprints. However, it is unclear whether this trend is reflective of other media outlets (e.g., those outside of Germany) and whether there are disciplinary differences in coverage trends.

More broadly, although preprints made up a significant proportion of the COVID-19-related literature available within the first months of the pandemic, it is unclear how media coverage of preprints compares to coverage of peer reviewed research. One article found that the five COVID-19-related research articles that received the most media coverage were all peer reviewed publications; however, the analysis was descriptive and did not compare the volume of preprint coverage to that of peer reviewed papers (Kousha & Thelwall, 2020). Another small study found no significant difference in the amount of media coverage received by medRxiv preprints and peer reviewed publications about COVID-19-related therapies that were posted between February 1–May 10, 2020 (Jung et al., 2021). Besançon et al. (2021) used Altmetric to examine news coverage of COVID-19-related preprints posted to arXiv, medRxiv, and bioRxiv between January–July 2020, finding that these preprints received more coverage than the non-COVID-19-related preprints posted to arXiv during the same time period. Again, coverage of preprints before the pandemic period was not considered. Fraser et al. (2020) found that bioRxiv preprints submitted between November 2013 and December 2017 received far less media coverage than either their peer reviewed versions or a control set of peer reviewed articles that were never deposited to bioRxiv. Finally, Waltman et al. (2021) found that, although some COVID-19-related preprints were highly reported on, overall, news coverage of peer reviewed literature outstripped coverage of preprints. Unfortunately, Waltman et al. (2021) did not report the average attention received per preprint vs peer reviewed article. However, the authors did examine news coverage received by a sample of high-profile preprints and their corresponding peer reviewed articles. For 45% of these preprint-article pairs, the preprint received more than 20% of the total news attention; for 11% of the pairs, preprints received more than 80% of the coverage (Waltman et al., 2021). Again, the authors did not compare these findings to rates of coverage before the pandemic.

Collectively, these results provide some of the first evidence that preprints have historically received less media coverage than peer reviewed research and that this trend may have started to shift during the pandemic. However, given the mixed and incomplete body of evidence, several questions remain unanswered. In particular, it is unclear whether the volume of preprint media coverage increased, decreased, or remained relatively stable in the years leading up the pandemic—information that could help shed light on whether preprint-based media coverage is likely to continue post-COVID-19. It is also unclear whether any changes in coverage seen during the pandemic apply only to COVID-19-related preprints or reflect a change in journalists’ willingness to use preprints in general. As such, to examine whether the pandemic has truly introduced a “paradigm shift” in journalistic practice, this study uses a sample of preprints that received coverage in English-language media between 2014–2021 to examine the following research questions:

> RQ1: Has the share of preprint coverage in the media increased during the COVID-19 pandemic?

> RQ2: Do changes in media coverage of COVID-19-related preprints extend to coverage of preprints on other topics?

### 2.2 Preprint media coverage in an evolving media landscape

It is also unclear from previous research which types of media outlets have driven media coverage of preprints and whether this has changed as a result of the pandemic. Journalism has evolved in important ways in the years leading up to the COVID-19 crisis, with financial pressures, shrinking news audiences, and changes to the digital communication landscape contributing to declines in specialized science journalism around the world (Saari et al., 1998; Schäfer, 2017). These declines have likely influenced the amount of media coverage that research articles—including preprints—receive, as outlets specializing in science appear to cover more research than general interest publications (Wihbey, 2017). In addition, an array of actors who have historically been considered “peripheral,” or outside of journalism, have entered the field, including bloggers, news aggregators, and other alternative outlets (Hermida, 2019; Schapals, 2022; Stocking, 2019). These peripheral actors may not always adhere to the established norms and practices that shape media coverage at traditional—or “legacy”—outlets (e.g., Harrison et al., 2020; Hurley & Tewksbury, 2012), which may affect how or whether they cover preprints. For example, journalists working at peripheral outlets may not be expected to adhere to professional journalism resources, such as the AP Style Guide, which recommend avoiding research that has not been peer reviewed (Froke et al., 2020; Haelle, 2020). Yet, both peripheral and legacy outlets actively covered COVID-19-related preprints during the early months of the pandemic (Fleerackers et al., 2021). Similarly, outlets which publish content but are not considered journalism, such as university websites and press release distribution services, may also contribute to mobilizing preprint research. For example, the Science Media Centre in Germany—a non-journalistic outlet that provides science journalists with access to research and expert perspectives—began sharing roundups of newly posted preprints during the pandemic (Broer, 2020; Broer & Pröschel, 2022). Again, however, any evidence about the nature of non-journalistic outlets reporting on preprints is limited to the pandemic period. As such, our third research question asks:

> RQ3: Have changes in media coverage of preprints occurred similarly across media outlets?

### 3.0 Method and materials

To identify media coverage, this study relies on data from Altmetric,^1^ a company that tracks mentions of research outputs across a range of digital media, including news media. Research suggests that Altmetric’s “Mainstream Media” category is a relatively reliable source of data but only when working with a predefined list of English-language media outlets (Fleerackers et al., 2022a; Ortega, 2020b, 2020a). In addition, because Altmetric regularly updates both the list of media outlets^2^ and research outputs^3^ it tracks, the volume of media coverage it collects may vary over time in ways that are unrelated to actual changes in news reporting. For these reasons, we decided to gather two datasets:

1. A primary dataset comprising news mentions of bioRxiv, medRxiv, arXiv, and SSRN preprints;
2. A comparison dataset comprising news mentions of peer reviewed research indexed in the Web of Science (WoS).

### 3.1 Identifying and characterizing media outlets that frequently cover research

Data were queried from local snapshots of the Web of Science and Altmetric Databases housed at the Observatoire des sciences et des technologies (OST).^4^ Data filtering and cleaning were performed using the Python pandas package (The Pandas Development Team, 2023). To identify our predefined set of media outlets, we queried a snapshot of the Altmetric database from June 3, 2021, for news mentions of all WoS research outputs associated with a digital object identifier (DOI). We restricted our search to mentions of research outputs that had been published in 2013 or later and that were mentioned in news stories between January 1, 2014, and June 3, 2021. We then filtered for outlets that consistently covered a high volume of research, defined for the purposes of this study as outlets that mentioned at least 100 WoS research items per year from 2014–2020. We manually checked the resulting 128 media outlets by visiting the URLs for their home pages provided by Altmetric. After excluding 25 outlets that were not written in English, five that were not tracked by Altmetric from 2021–2022 (e.g., because they had changed their domain names), three whose URLs did not resolve, and one with all misidentified mentions, we were left with a final sample of 94 outlets.

Next, we applied a coding protocol adapted from Hermida & Young (2019) to characterize the nature of these media outlets. We analyzed each outlet’s main topical focus (e.g., science and technology, health and medicine, general news, etc.) and assessed whether it was best described as legacy journalism (i.e., staffed by professional journalists who adhere to traditional journalistic norms), peripheral journalism (i.e., staffed by individuals who have traditionally worked outside of journalism and who adhere to emerging or alternative norms), or non-journalism (i.e., organizations such as universities, press release services, or academic journals that do not produce journalism). A detailed version of the coding protocol, including examples, is available from Fleerackers and Fagan (2022).

Coding was performed by researchers with professional journalism experience: the lead author and a research assistant who was not aware of the study objectives (cf. Hermida & Young, 2019). The two coders independently explored the media outlets’ websites, examining their content, Mission Statement, and, if available, other relevant pages (e.g., Masthead, Editorial Guidelines, Code of Conduct). The coders compared their coding and resolved any discrepancies through discussion, and, if needed, by consulting an outside researcher (also a former journalist). Such double coding approaches are appropriate when data are not very numerous (Krippendorff, 2004), as in the present study. Results of the final coding are reported in aggregate in Table 1; coding for the full list of outlets is available at (Alperin, Fleerackers, et al., 2023).

**Table 1.** Nature of media outlets that frequently cover Web of Science research

| Topic | N | Type | N |
| --- | --- | --- | --- |
| General News | 38 | Legacy | 49 |
| Science/Technology | 28 | Peripheral | 35 |
| Health/Medicine | 15 | Non-journalism | 10 |
| Other | 13 |  |  |
| Total | 94 |  | 94 |

### 3.2 Gathering news mentions of preprint research

We queried Altmetric for mentions of bioRxiv, medRxiv, arXiv, and SSRN preprints in stories published by the 94 outlets since January 1, 2014. This yielded a total of 40,039 mentions of 15,041 preprints across 31,258 news stories. For each of these preprints, we gathered the publication dates from the arXiv and Crossref APIs using the Python arxiv and habanero packages (Chamberlain, 2020; Schwab, 2021).

Next, because previous research suggests publication date metadata can often be incorrect or incomplete (Haustein et al., 2015), we manually checked subsamples of our data and compared the publication dates provided by Crossref, the arXiv API, and Altmetric. The most reliable publication date for each server was retained for analysis. For bioRxiv and medRxiv, this was the DOI creation date (i.e., the date that the DOI for the preprint was deposited in Crossref); for arXiv, it was the date provided by the arXiv API; and for SSRN, it was either the “first posted on” date provided by Altmetric or Crossref’s DOI creation date, whichever came first. We removed 3,619 preprints that were published before 2013, as these publication dates were particularly unreliable (perhaps because Altmetric started tracking mentions partway through 2012 and thus has incomplete data for previously published outputs)^5^. Even after excluding these preprints and selecting the most reliable publication date for each server, we noted that publication dates for *arXiv* and *SSRN* sometimes differed from the dates visible on the server web page by a few days—a limitation that we kept in mind during data cleaning and analysis.

We made several further exclusions to ensure that the mentions in our dataset were mentions of true preprints (i.e., rather than postprints or published versions of preprints). First, we removed 165 mentions of postprints, which we defined as preprints that were posted on the same day, or after, their published versions were published. Because, as mentioned above, publication dates for preprints were often incorrect by a few days, we excluded an additional 332 mentions of preprints with a publication date within seven days of the published version’s publication date (i.e., suspected postprints). We also removed 327 mentions of preprints in news stories that were published before the preprint was first posted, using a five-day cut off to allow for the slight inconsistencies we identified in the publication metadata. Because Altmetric does not disambiguate between preprints and published versions for some preprint servers^6,7^ and may thus erroneously include some mentions of peer reviewed research, we removed 3,547 mentions in news stories published after the peer reviewed version of the preprint was published, again using a five-day margin. While this approach may have removed some true mentions of preprints, these false removals are likely limited, as it is relatively uncommon for news stories to mention research outputs more than a few weeks after initial publication (Maggio et al., 2017). Finally, we removed an additional 1,021 duplicate news mentions (where the same preprint was mentioned in the same story more than once). In total, filtering led to the exclusion of 9,081 mentions (22.5% of the original dataset). The code used for filtering has been made publicly available (Alperin, Shores, et al., 2023). Our final preprint sample comprised 31,028 mentions of 11,538 preprints by the 94 outlets in our sample (Alperin, Fleerackers, et al., 2023).

### 3.3 Gathering news mentions of peer-reviewed research

We downloaded all the mentions of WoS research from our 94 outlets, resulting in 1,657,202 mentions of 466,138 distinct research outputs. From these, we filtered 156,187 mentions of research articles that were published prior to 2013, 579 mentions that were already included in the preprint data, and 14,482 duplicate mentions (where an article was mentioned in the same news story more than once). In total, filtering led to the exclusion of 170,669 mentions (10.3% of original dataset).

The final published research sample comprised 1,486,533 mentions of 397,446 distinct peer reviewed research outputs by the 94 outlets (Alperin, Fleerackers, et al., 2023).

### 3.4 Identifying news mentions of COVID-19 research

To identify COVID-19-related preprints and WoS outputs, we searched for the presence of the following COVID-19-related keywords in the outputs’ titles using R version 4.3.0 (2023): *coronavirus, covid-19, sars-cov, sars-cov-2, ncov-2019, 2019-ncov, hcov-19, sars-2, pandemic, covid, Severe Acute Respiratory Syndrome Coronavirus 2, 2019 ncov*. These keywords were a combination of those used by Fraser et al. (2021) and those listed in the National Library of Medicine’s search strategy for identifying COVID-19-related literature (Chen et al., 2020). We also added the term “pandemic,” which wasn’t included in either of these lists of keywords but is likely used in many COVID-19 titles. As some keywords (e.g., “pandemic”) may have been used in non-COVID-19 contexts, we also filtered for research published in 2020 or later when identifying COVID-19-related research.

### 3.5 Statistical analyses

Statistical analysis was performed using Stata version 17 (StataCorp, 2021). The Stata script used for the following analysis has been made publicly available (Alperin, Shores, et al., 2023). Throughout our analyses, we examined changes in preprint media coverage in terms of proportions, rather than counts. Specifically, we compared mentions of preprints against mentions of all research in our sample (i.e., mentions of preprints *and* WoS research). Doing so allowed us to control for any fluctuations in the volume of preprint mentions that were created by changes in Altmetric’s approach to identifying research mentions during the study period, rather than the result of changing journalistic practices. For ease of reading, we use the term “share of preprint mentions” to refer to the proportion of all research mentions that focused on preprints and “share of WoS mentions” to refer to the proportion that focused on WoS research.

To answer RQ1, we created a model (Equation (1)) to estimate the degree to which medRxiv and COVID-19 contributed to changes in the volume of media coverage of preprints after 2019. Disentangling any change in preprint coverage due to the launch of the server and the onset of the pandemic was necessary as the creation of medRxiv preprints in 2019 (Kaiser, 2019) coincided closely with the start of the COVID-19 era. As such, in Equation (1), we regressed a binary indicator coded as 1 if the media mention referenced a preprint and coded as 0 otherwise against time, encoded as linear days since Jan 1, 2014 and allowed to be identified with 3rd-order polynomial trends (β_1_through β_3_), with each vector of 3rd-order polynomial terms estimated in both the pre-COVID-19 era and COVID-19 era (α_0_ interacted with the vector of time trends). We differentiated pre-COVID-19 from COVID-19 era mentions through a binary indicator, coded as 1 if the preprint was mentioned in a media story published after January 10, 2020 (i.e., when the WHO first used the term “2019-nCoV” to describe the novel coronavirus; WHO, 2020), and coded as 0 otherwise. We modeled the period between the first media mention of a medRxiv preprint (i.e., on July 23, 2019, which postdates the launch of the site on June 25, 2019 by about one month) and the WHO’s statement as a linear intercept shift (β_4_). In practice, this variable allowed us to differentiate the change in preprint mentions that occurred with the introduction of medRxiv before (but close to the onset of) COVID-19 from the effect of COVID-19 itself. Similarly, we modeled the mentions of preprints with titles that included COVID-19-related language (i.e., “sars-cov-2” or a related term) as a linear intercept shift (β_5_). This last variable is important, as it allowed us to differentiate the change in preprint mentions for COVID-19-related topics in the media from changes in preprint prints in the COVID-19-era but not about COVID-19 topics. Lastly, to adjust for seasonality and periodicity effects we controlled for week-of-year intercepts (γ_*wy*_; e.g., first week of 2014) and day-of-month effects (δ_*md*_; e.g., Tuesdays in January). In practice, controlling for periodicity and seasonality had little effect on model parameters but allowed us to rule out correlations between period effects and the onset of COVID-19.

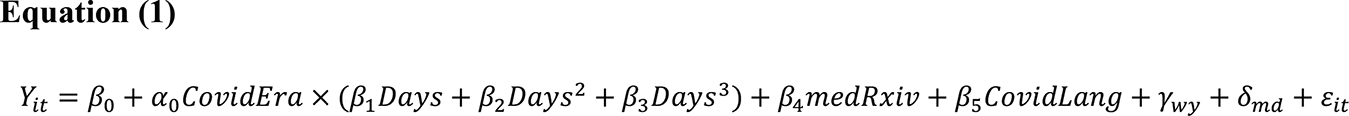

Next, we estimated separate regressions that allow us to test whether changes in preprint mentions vary across (a) preprint servers, (b) media outlets focused on different topics, and (c) media outlets of different types. Because preprint servers necessarily represent preprint mentions, we discarded mentions of articles from WoS and collapsed the data so that we could observe counts of preprint mentions by day and identify any changes in these counts among the four servers (RQ2). To identify changes among the four media outlet topics and three outlet types, respectively, we kept the data as described previously, with each row representing a unique media mention of a preprint or WoS article. To identify changes in the share of preprint mentions across the four media outlet topics (RQ3), we focused on the three most prevalent topics in our sample—Health/Medicine, General News, and Science/Technology—and an “Other” category that included a variety of other topics (e.g., Business, Lifestyle, Explicit Point-of-View).

Because we are exploring heterogeneity across servers, topics, and types, we simplified the regression Equation (1) by replacing the linear, quadratic, and cubic *Days* variables with month-by-year fixed effects (λ_*my*_ in Equation (2) below). These fixed effects control for time trends non-parametrically in a similar way as in Equation (1) but without the need to directly identify the time effects (i.e., these time effects are partialled from the regression equation as “nuisance parameters”). Our estimation equation for these heterogeneous changes therefore appears as follows:

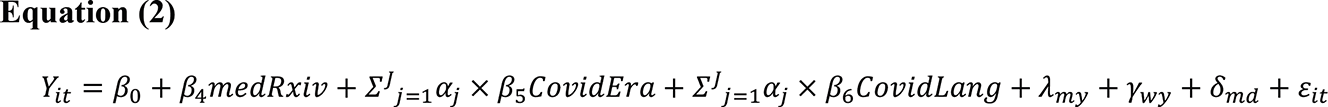

where the key difference is that we identify changes in the pandemic era without and with COVID-19-related titles (α_&_ × β_)_and α_&_ × β_*_, respectively) for the four preprint servers (j=1 through J=4), the four outlet topics (j=1 through J=4), and three outlet types (j=1 through J=3).

## 4.0 Results

### 4.1 Has the share of preprint coverage in the media increased during the COVID-19 pandemic?

Our models suggest that the annual number and share of preprint mentions increased slowly from 2014–2019, then increased dramatically in 2020-2021 (Table 2, Figure 1). However, even during the pandemic period, preprint mentions made up only a small subset of media coverage of research, at less than 5% of all mentions of research. We also saw evidence of a shift in which servers received the most attention during the pandemic. Before the introduction of COVID-19, most mentions of preprints cited preprints posted to arXiv or SSRN; yet during the pandemic, bioRxiv and medRxiv became the most frequently mentioned servers.

**Table 2.** Number and share of preprint mentions

| Year | Number of<br>WoS<br>mentions | Number of<br>preprint<br>mentions | arXiv<br>preprints | SSRN<br>preprints | bioRxiv<br>preprints | medRxiv<br>preprints | Total<br>preprint<br>proportion |
| --- | --- | --- | --- | --- | --- | --- | --- |
| 2014 | 98,580 | 753 | 0.34% | 0.41% | 0.01% | N/A | 0.76% |
| 2015 | 122,641 | 1,263 | 0.42% | 0.56% | 0.04% | N/A | 1.02% |
| 2016 | 192,414 | 2,115 | 0.44% | 0.56% | 0.09% | N/A | 1.09% |
| 2017 | 214,714 | 2,412 | 0.42% | 0.60% | 0.09% | N/A | 1.11% |
| 2018 | 218,946 | 2,597 | 0.48% | 0.58% | 0.12% | N/A | 1.17% |
| 2019 | 231,503 | 3,275 | 0.55% | 0.67% | 0.16% | 0.01% | 1.39% |
| 2020 | 279,737 | 12,484 | 0.46% | 0.56% | 0.85% | 2.40% | 4.27% |
| 2021* | 127,998 | 6,129 | 0.50% | 0.29% | 1.27% | 2.51% | 4.57% |
| Total | 1,486,533 | 31,028 |  |  |  |  |  |
| % of all<br>research<br>mentions |  |  | 0.46% | 0.55% | 0.35% | 0.68% | 2.04% |
| % of all<br>preprint<br>mentions |  |  | 22.62% | 26.93% | 16.95% | 33.49% | 100% |

\* partial year

**Figure 1.**
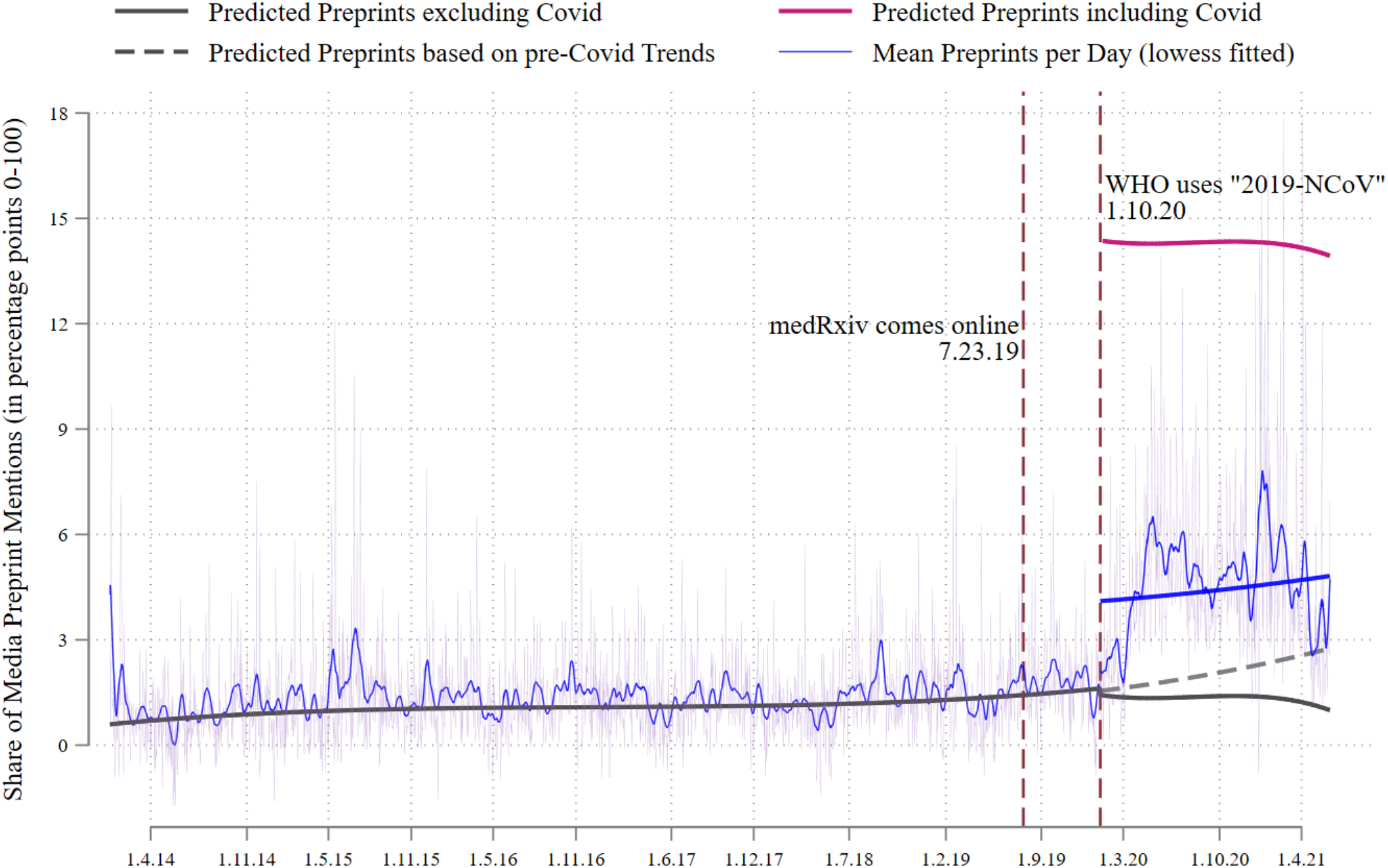
Share of preprint mentions per day. NB: Figure shows the average proportion of preprint mentions in the media per day beginning Jan 1, 2014, and ending June 3, 2021. The fluctuating faded purple line plots residualized mean share of preprint mentions per day, controlling for week-of-year effects (e.g., first week of 2014, second week of 2014, etc.) and day-by-month effects (e.g., Tuesdays in January); the fluctuating blue line is the local linear regression (lowess) fitted line of those data. The solid gray line is the predicted share of daily preprint mentions, with prediction based on a 3rd-order polynomial function, controlling for week-of-year effects and day-of-month effects. We estimate an intercept shift for the period before COVID-19 and after medRxiv was introduced on July 23, 2019, and an intercept shift for mentions of preprints with titles that include COVID-19-related language (e.g., “novel coronavirus” or a related term). The solid fuchsia line represents this last effect and is the estimated change in mean preprint mentions per day among all COVID-19-related mentions of research (preprints and WoS publications). The dashed gray line is the predicted share of daily preprint mentions based on trends observed in the period prior to the emergence of COVID-19. The solid blue line describes the proportion of preprint mentions in the COVID-19 era and is the weighted average of the fuchsia and gray lines. See Equation 1 for details.

With respect to medRxiv preprints, we found that the onset of COVID-19 increased the share of preprint mentions in the media, beyond any increase due to the launch of the server in 2019. Specifically, we estimate that, prior to the introduction of medRxiv and COVID-19, the share of preprints mentioned in the media was increasing at a glacial pace (an *annual* rate of 0.21 percentage points; p-value<0.000; 95% CI [0.13 - 0.29]; see solid gray line, Figure 1). When medRxiv was introduced, the share of preprint mentions did not change (estimated decrease=0.005 percentage points; p-value=0.957; 95% CI [-0.17 - 0.16]). In contrast, the share of preprint mentions increased by an estimated 2.58 percentage points after the onset of the pandemic (p-value<0.000; 95% CI [2.45 - 2.70]; see solid blue line). This significant but modest increase applied to all preprint mentions, but masks large differences in the proportion of preprint mentions between COVID-19-related and non-COVID-19-related research during the pandemic.

Indeed, our model strongly suggests that preprints played a far greater role in media coverage of COVID-19 specifically rather than in coverage of other topics. This can be seen from the “COVID-19” line (in fuchsia) in Figure 1, which represents the estimated share of preprint mentions among all the mentions of COVID-19-related research (i.e., both preprints and WoS articles that included COVID-19-related language in the title). We estimated an increase in these COVID-19-related preprint mentions of 12.94 percentage points (p-value<0.000; 95% CI [12.84 - 13.04]), a large increase relative to predicted preprint mentions based on pre-COVID-19 trends (gray dotted line). We explore coverage of non-COVID-19 preprints in more detail in Section 4.2.

We further tested whether any changes in the share of preprint mentions seen during the pandemic could be linked to changes in mentions of WoS research during this period. We implemented this test by comparing growth rates of media mentions for preprints and WoS research over time. Given that preprint mentions comprised only about 2 percent of all mentions in our sample and to place preprint and WoS mentions on a common y-axis, we plotted preprint and WoS mentions as growth rates. Growth rates for preprint and WoS mentions were each calculated using the total number of mentions in the first 28 days of our data beginning with Sunday (i.e., January 5, 2014). These mentions in the first 28 days comprised our “base rate,” and the total number of mentions in each sequential 28 days were then scaled by that base rate.

Here, we find that the rise in the share of preprint mentions that took place during the pandemic was not simply an artifact of a decrease in WoS mentions. As can be seen from Figure 2, WoS mentions increased by about 2.3 percentage points between May 2014 to September 2019, and this pace of growth remained relatively unchanged after COVID-19 began and started to garner media attention. In contrast, preprint mentions had increased by about 5.7-fold by the time of the WHO’s announcement about “2019-nCoV” in January 2020, but skyrocketed to a 30-fold increase at the height of the pandemic in May 2020. This figure thus shows that the increase in the proportion of preprint mentions during the pandemic era was driven almost entirely by an increased number of preprint mentions and not a decrease in the number of WoS mentions.

**Figure 2.**
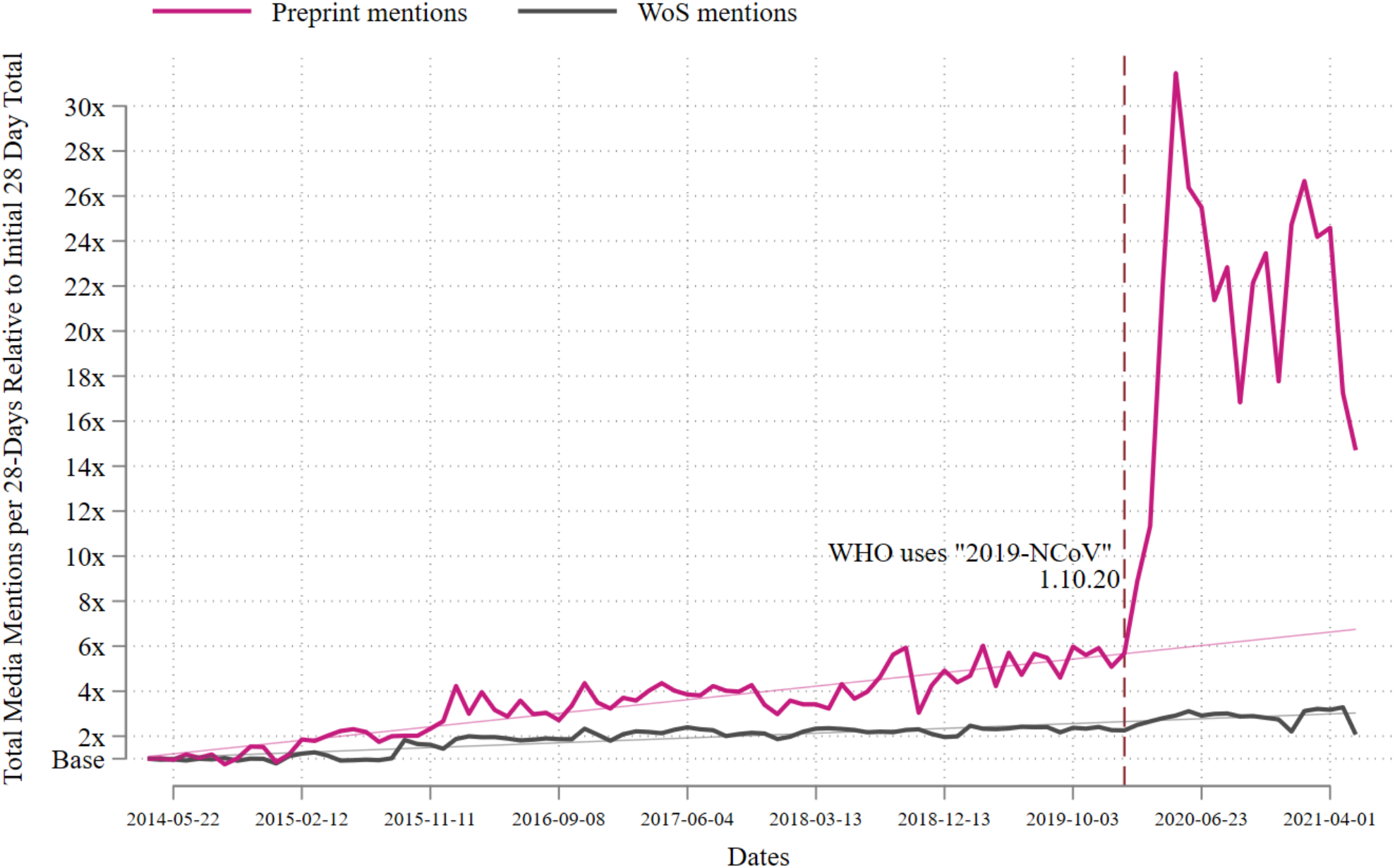
Growth rates of mentions for preprints and Web of Science (WoS) articles over time. NB: Figure shows the number of preprint and Web of Science (WoS) mentions in the media per 28 days beginning the first 28 days of March 2, 2014, and ending June 3, 2021. We select the first 28 days starting with March 2, 2014, for stylistic reasons, as preprint mentions declined slightly between January 2014 and March 2014; however, starting on January 1 does not change any of the findings we discuss below. Because preprint mentions only comprise about two percent of all media mentions in our data and to put these counts on a common y-axis, we scale monthly (28 day) preprint and WoS mentions by the number of mentions for preprints and WoS, respectively, by the initial number of mentions in the first 28 days of our sample beginning March 2, 2014. Thus, subsequent monthly mentions are relative to this base period. For example, a 28-day media mention count of “2x” means that media mentions in that 28-day period were two times larger than media mention counts from March 2, 2014 to March 29, 2014 (i.e., the initial 28 day period). The fuchsia and gray lines indicate 28-day preprint and WoS media mentions, respectively, in this relative metric.

### 4.2 Do changes in media coverage of COVID-19-related preprints extend to coverage of preprints on other topics?

Our results suggest that the onset of the pandemic not only increased media attention to COVID-19-related preprints but may have also decreased attention to preprints on other topics. Among all research that excluded COVID-19-related language (solid gray line, Figure 1), we found that the share of preprint mentions during the pandemic decreased by 0.18 percentage points, although this decrease was not significant (p-value=0.129; 95% CI [-0.42 - 0.05]). Model-based estimates suggest that by June 3, 2021, if the pandemic had not occurred, we would have expected the share of preprint mentions to be 2.58 percentage points (dashed gray line, Figure 1); yet the observed share of non-COVID-19-related preprint mentions comprised only 0.86 percentage points of all media mentions, a difference of 1.71 percentage points from what would have been expected (p-value<0.000; 95% CI [1.13 - 2.31]). This last result suggests that the pandemic may have shifted media attention away from preprints about non-COVID-19-related topics by modest amounts. In effect, our results suggest that COVID-19-related preprint mentions eclipsed pre-pandemic preprint mentions.

Looking at the number of preprint mentions by server, we observed that there was no increase in non-COVID-19-related preprint mentions in the pandemic for any server (Figure 3). All point estimates were trivially small—about 0.7 to 1.8 fewer mentions per day, on average—and not statistically significantly different from zero (p-values range from 0.217 to 0.699). For articles that included COVID-19-related language in the titles, there was an average increase in daily media mentions of bioRxiv and medRxiv preprints—of 6.2 (p-value<0.000; 95% CI [5.34 - 6.95]) and 19.2 (p-value<0.000; 95% CI [18.25-20.08]), respectively—and a significant decrease in average daily media mentions for arXiv and SSRN—of -2.7 (p-value<0.000; 95% CI [-3.75 – -1.68]) and -1.6 (p-value<0.000; 95% CI [-2.69 – -0.51]) mentions per day, on average. In total, for the 511 days in the pandemic era in our sample, this amounted to an increase of about 9,800 total mentions of medRxiv preprints and 3,170 total mentions of bioRxiv preprints.

**Figure 3:**
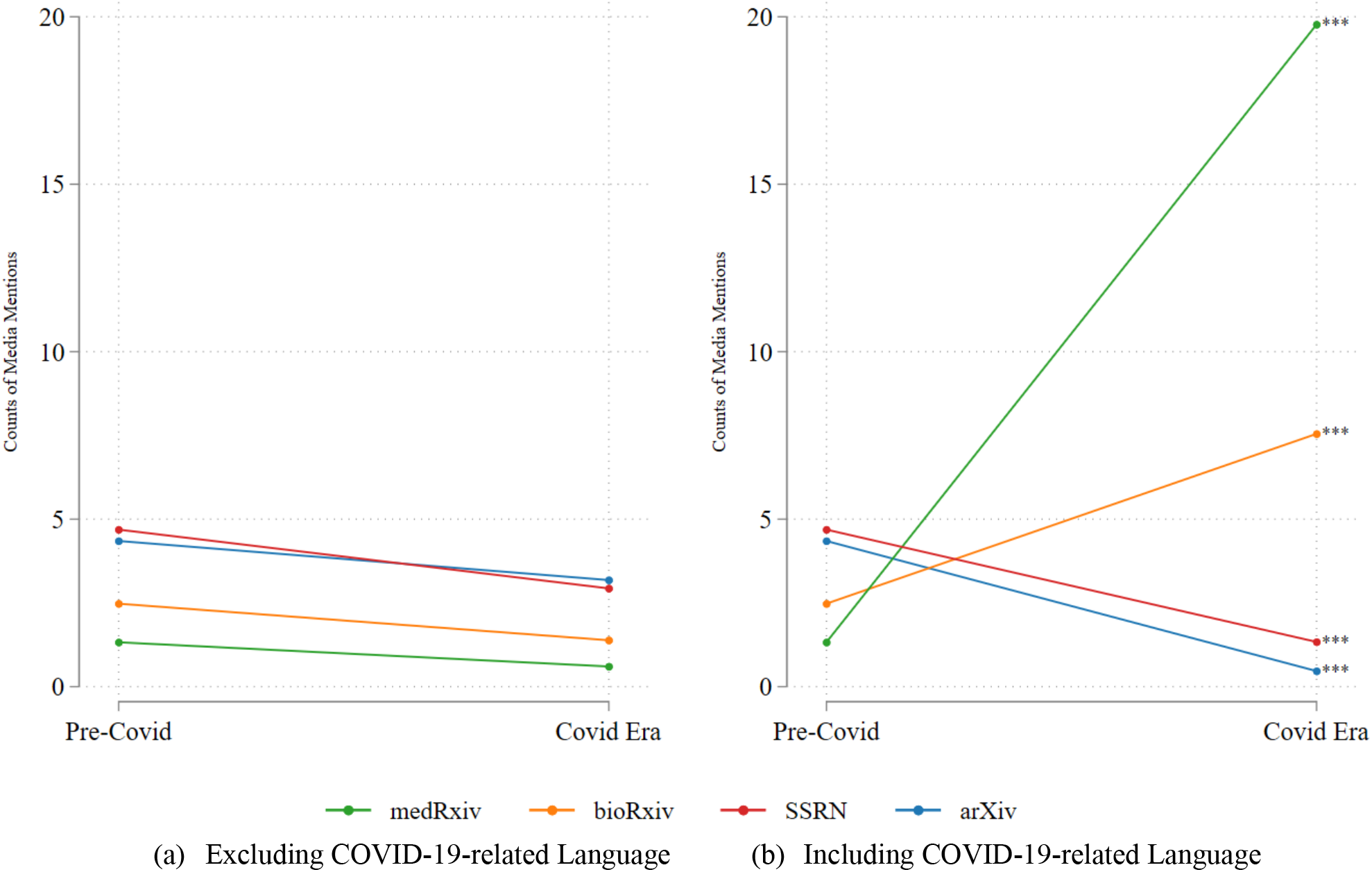
Change in the average daily count of preprint mentions, by server. NB: Model parameters are from Equations (2). Statistical significance is determined based on the test of whether pandemic era counts are different from pre-pandemic counts for mentions of preprints with titles that exclude COVID-19-related language (panel a) and include COVID-19-related language (panel b). Key: * 10% significance; ** 5% significance; *** 1% significance

It is important to note that the declines in mentions of arXiv and SSRN preprints were only significant for preprints that included a COVID-19-related keyword in the title. That is, the media were less likely to mention preprints from these servers that were about COVID-19; instead, when communicating about pandemic research, they tended to mention bioRxiv or medRxiv preprints. These results suggest that the media drew on the servers they expected would house the research most relevant to their area of interest. It also suggests that COVID-19-related coverage tended to focus on medical aspects of the pandemic and less so on social or economic aspects.

### 4.3 Have changes in media coverage of preprints occurred similarly across media outlets?

Finally, we tested how preprint mentions changed across media outlets with four different topical foci (i.e., General News, Science/Technology, Health/Medicine, Other) or of different types (i.e., legacy, peripheral, or non-journalism). For mentions of COVID-19-related research, we found that outlets in all four topic categories increased their preprint coverage dramatically during the pandemic, but to different extents. Increases ranged from 8.3 percentage points (Science/Technology) to 15.6 percentage points (Health/Medicine) and were all statistically significant (p-value<0.000 for all coefficients) (Figure 4). Changes in the share of mentions for non-COVID-19-related preprints were trivial, with only the “Other” category seeing a small but statistically significant increase (0.9 percentage points).

**Figure 4.**
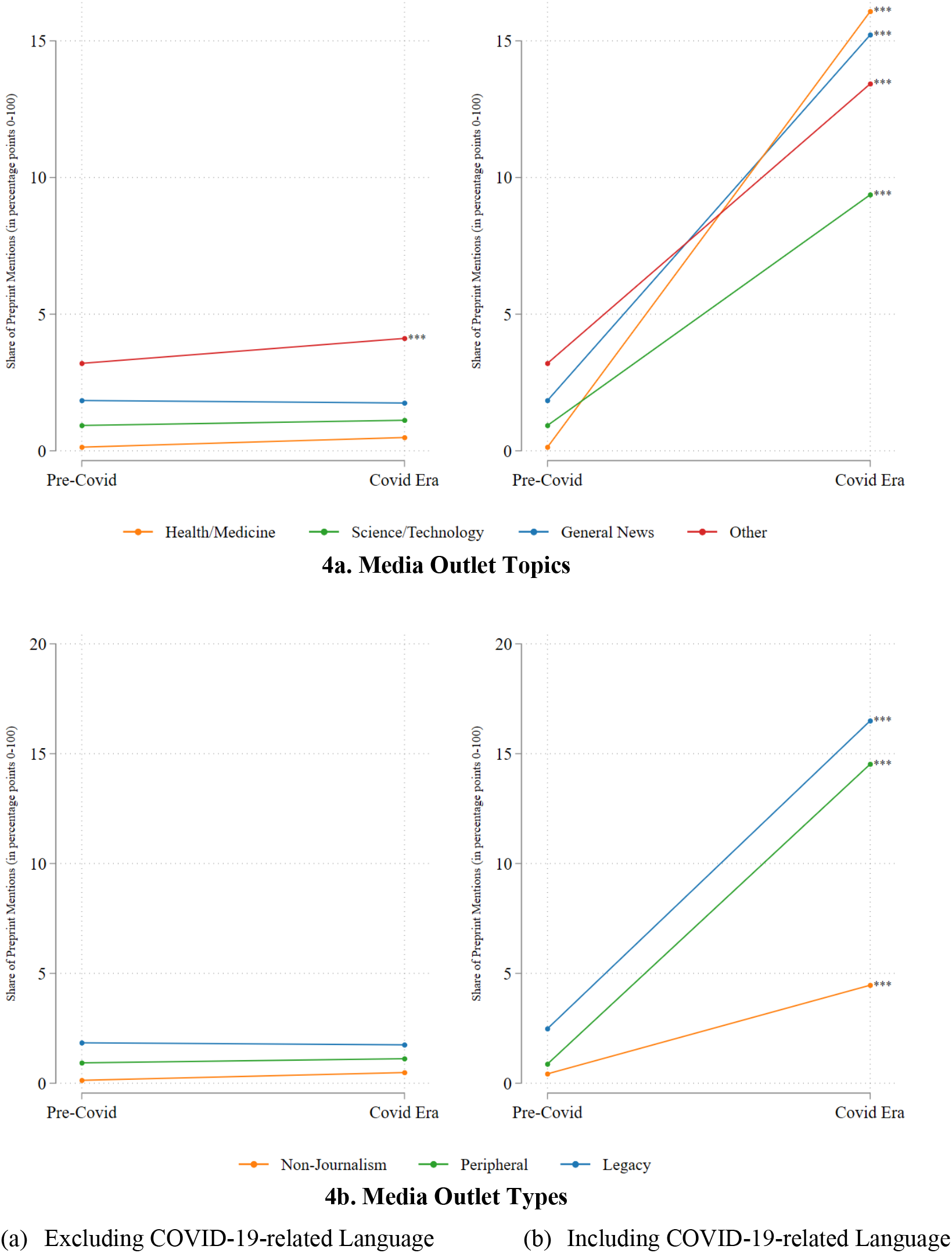
Change in the average daily share of preprint mentions, by topic and outlet. NB: Model parameters are from Equations (2). Statistical significance was determined by testing whether pre-pandemic share of preprint media mentions was different from those during the pandemic, both among mentions of research with titles that excluded COVID-19-related language (panel A) and those that included COVID-19-related language (panel B). Key: * 10% significance; ** 5% significance; *** 1% significance

Similarly, none of the outlet types (i.e., legacy, peripheral, non-journalism) saw a statistically significant increase in the share of preprint mentions of non-COVID-19-related articles, and the coefficients themselves were trivially small, never reaching 1 percentage point. However, just as with the topic-based data, all three outlet types increased the share of mentions of COVID-19-related preprints after the WHO announcement in 2020 (estimates ranged from 4 to 14 percentage points for non-journalism and legacy outlet types, respectively.

Finally, to provide a better sense of the nature of the outlets that frequently rely on preprints, we identified the 25 media outlets whose coverage included the largest share of research mentions in general (i.e., mentions of preprints and WoS outputs) and calculated their share of preprint mentions both before and during the COVID-19 era (Table 6). The list represents about 75% of all research mentions in our sample and includes a mix of legacy media, such as *BBC News* and *The New York Times*, and peripheral outlets, such as *Reason* or *The Conversation*. Several non-journalism outlets also appear on the list, mostly services such as *EurekAlert!* and *Newswise,* which do not publish original articles but distribute science press releases (many of which include mentions of new research). Among outlets that tended to cover a high proportion of preprints in general, the US libertarian magazine *Reason* stood out, mentioning approximately one preprint for every three WoS outputs—far more than any other outlet in our sample prior to the COVID-19 pandemic. Interestingly, the outlet’s share of mentions actually decreased slightly during the pandemic, from 27% to 24%. Among the outlets that saw the largest increase in their share of preprint mentions, the peripheral Health/Medicine outlet *News Medical* topped the list, with essentially no preprint mentions before the pandemic but a share of 43% during the pandemic. Several major legacy General News outlets, such as *BBC News, The Daily Mail, The New York Times,* and *The Guardian,* also saw notable increases in preprint coverage, moving from minimal use of preprints to covering about one preprint for every four or five mentions of research. Although some specialized Science/Technology outlets (e.g., *Scientific American, Phys.org*) increased their coverage of preprints during COVID-19, these increases tended to be less pronounced than those seen among the major General News outlets.

**Table 6.** Largest 25 media outlets based on mentions and the share of mentions that include preprints

|  | Outlet's Share of All<br>Research Mentions | Outlet's Share of Preprint Mentions |  |
| --- | --- | --- | --- |
|  |  | Pre-pandemic Era | Pandemic Era |
| BBC News | 1.75% | 1.69% | 21.85% |
| Business Insider | 2.51% | 2.61% | 17.45% |
| Business Insider Australia | 1.37% | 2.03% | 16.32% |
| Daily Mail | 2.07% | 1.45% | 22.24% |
| EurekAlert! | 1.55% | 0.24% | 2.83% |
| Forbes | 6.37% | 9.94% | 12.37% |
| Gizmodo | 1.00% | 3.33% | 14.83% |
| MedicalXpress | 2.42% | 0.19% | 7.75% |
| New Scientist | 1.09% | 4.15% | 20.84% |
| New York Times | 8.12% | 3.69% | 23.32% |
| Newswise | 1.28% | 0.60% | 6.98% |
| Phys.org | 3.86% | 0.99% | 5.63% |
| Quartz | 1.47% | 4.93% | 16.38% |
| Reason | 2.03% | 26.83% | 24.01% |
| Salon | 1.25% | 4.27% | 17.27% |
| Science/AAAS | 0.98% | 2.85% | 33.62% |
| Scientific American | 1.02% | 2.13% | 17.17% |
| The Atlantic | 1.57% | 6.48% | 23.82% |
| The Conversation | 6.84% | 1.75% | 9.76% |
| The Guardian | 1.88% | 2.01% | 20.56% |
| Medical News | 8.73% | 0.30% | 42.70% |
| Times of India | 1.03% | 0.99% | 11.96% |
| Vox | 1.60% | 4.79% | 19.23% |
| Washington Post | 3.16% | 4.66% | 13.68% |
| Yahoo! News | 11.44% | 1.57% | 14.27% |

NB: Top 25 outlets shown based on their share of mentions of Web of Science outputs and preprints, representing about 75% of all mentions in our sample. Column 2 (Outlet's Share of All Research Mentions) shows each outlet's share of all mentions of Web of Science outputs and preprints. Columns 3 and 4 (Outlet's Share of Preprint Mentions) shows the share of each outlet's mentions of preprints prior to COVID-19 and the share of each outlet's mentions for preprints during COVID-19. Grayscale conditional formatting is based on column 2 alone and then columns 3 and 4 jointly.

## 5.0 Discussion

It has been argued that preprint coverage during the pandemic constituted a break from journalism norms and a paradigm shift in how emergent research is reported on and shared with the public (Burke, 2021; Makri, 2021). Using longitudinal data from the Web of Science (WoS) and four preprint servers, this study sought to establish whether, in what ways, and to what extent this is the case. By identifying how the volume and nature of preprint media coverage has changed over time and what role the pandemic has played in this change, our study makes an important contribution to our understanding of journalists’ use of preprints—a topic about which much has been written, but very little is actually known.

A key finding from our analysis is that the volume of preprint media coverage increased by roughly fourfold in the pandemic period, a clear break from the slight but steady upward trend that preceded it. Virtually all of this increase was driven by coverage of COVID-19-related preprints, with little change in coverage of preprints on other topics. Although coverage of peer reviewed research continued to exceed preprint coverage—even during the height of the crisis—the growth in coverage of preprints seen during this period may imply a shift in journalistic norms and practices, including a changing outlook on preliminary, unvetted research and its reporting.

At the same time, however, we observed a slight (but nonsignificant) decrease in coverage of non-COVID-19-related preprints during the pandemic. This lack of coverage of non-COVID-19-related preprints may simply be the result of outsized media attention to COVID-19 in general (i.e., not just COVID-19-related preprints), which may have come at the expense of coverage on other topics. Yet, it could also indicate that the surge in preprint coverage observed during the pandemic was a temporary change—a break from established norms that journalists made to cover a rapidly evolving crisis, rather than a true shift in practice. More research is needed to assess the degree to which increases in preprint coverage will persist in the coming years, as media outlets and scientists turn their attention away from COVID-19 and toward other issues.

Interestingly, the sharp rise in preprint coverage seen during the pandemic was most pronounced for health and medical outlets, which appear to have been resistant to covering preprints until relatively recently. While outlets specializing in other topics, such as science and technology, covered preprints at least occasionally before the pandemic, our findings suggest that, for health and medical outlets, the crisis seems to have created something closer to the “paradigm shift” described by journalists in previous research (Fleerackers et al., 2022a). Preprints were barely mentioned in health and medical outlets up until 2019—even after medRxiv was launched—but become a frequent source of coverage in these outlets after 2020, particularly when reporting on COVID-19. Again, more research is needed to assess whether this trend will continue beyond the pandemic.

The factors that motivated health and medical journalists to adapt their practices during COVID-19 also remain unclear. While the medical nature of the crisis likely played a primary role, at least some of this shift may be linked to a parallel shift in preprint use among health and medical scholars themselves. Like journalists, researchers in these areas have historically been hesitant to post or cite unreviewed research (Flanagin et al., 2020; Maslove, 2018), but became active users of preprints during the pandemic (Fraser et al., 2021; Waltman et al., 2021). Since journalists who report on research rely heavily on interviews with scientific experts (Schultz, 2023), changing attitudes toward preprints among medical scientists would likely affect reporting practices on medical and related issues. It is possible, in other words, that the uptake of preprints by medical and health outlets reflects the growing acceptance of preprints within the medical and health sciences. This may also be true of preprint-based journalism more broadly, as preprint adoption also grew during the study period. Waltman et al. (2021) report that the number of preprints in 2020 was about 150% larger than the number of preprints in 2015, while Penfold and Polka (2020)—working with data from PubMed and 10 preprint servers—found that the number of biology preprints increased almost tenfold between 2013 and 2019 (from 0.24% to 2.36%). However, just because more preprints are becoming available doesn’t mean journalists will automatically cover them. By covering preprint science, journalists may—potentially—be adapting their own norms to follow those of scientists.

In terms of outlet types, we found that both traditional, legacy outlets (e.g., *The New York Times*) and peripheral media outlets (e.g., *News Medical*) were covering preprints to some extent before the pandemic, but greatly increased this coverage during the crisis. The similar pattern seen for the two outlet types is surprising, as peripheral media outlets are often conceptualized as following different norms, ethics, and practices than legacy media and as being less beholden to professional guidelines, such as those that urge journalists to avoid covering unreviewed research (Froke et al., 2020). Our findings thus align with previous scholarship which has suggested that the boundaries between legacy and peripheral journalism are blurring and that categorizing outlets this way may no longer be meaningful (Deuze & Witschge, 2018; Witschge et al., 2019). While more research is needed, it is possible that such blurring boundaries are especially likely in contexts where professional norms are not yet well-established, such as when reporting on preprints. Future studies could explore whether the similarities we observed in preprint coverage among peripheral and legacy outlets also apply to larger and more diverse outlet samples, or to other situations in which journalistic practices are rapidly evolving.

Collectively, our findings provide some of the first evidence that journalists are increasingly using preprints—at least in some areas—and that the pandemic has greatly accelerated this use. However, this conclusion should be considered alongside several limitations. First, there are known challenges of working with Altmetric data to identify media coverage of research, particularly in languages other than English (see Ortega, 2020a, for a review). We have attempted to mitigate these challenges by working with a predefined set of English-language media outlets, as recommended in previous research (Fleerackers et al., 2022b). Yet, while the restricted nature of our sample of outlets is a strength of this study, it is also a limitation, as the patterns we observed among these 94 outlets may not apply to those that less frequently report on research or do so in different languages. Replicating our findings with a larger set of outlets or through complementary data gathering methods would be a fruitful avenue for future research.

We also aimed to make our findings more robust by contextualizing any increases in preprint media coverage alongside changes in coverage of peer reviewed research during the same time period. To do so, we relied on Web of Science data, which is biased towards studies from scientific, technical, and medical disciplines and published in English-language journals from the Global North (Alperin et al., 2014; Mongeon & Paul-Hus, 2016). Given our study’s focus on English-language media outlets, the impact of the language bias is likely minimal (i.e., it is relatively unlikely that a journalist working for an English-language outlet would cover non-English research). However, the disciplinary and geographic biases are limitations of our study that should be kept in mind when interpreting the results.

Finally, the nature of our data only enabled us to explore changes in preprint media coverage from 2014 through the first year and a half of the pandemic, leaving many questions unanswered about what the future will bring. We hope that scholars will build on our findings to provide further insight into the implications of the preprint coverage seen during COVID-19 will persist long-term.

## Acknowledgements

The authors wish to thank Altmetric for providing this study’s data free of charge for research purposes and Stefanie Haustein for her input on the methodology.

## Footnotes

1 altmetric.com

2 https://help.altmetric.com/support/solutions/articles/6000235999-news-and-mainstream-media

3 https://www.altmetric.com/about-our-data/how-it-works-2/

4 https://www.ost.uqam.ca/

5 NISO Altmetrics Working Group C “Data Quality” ‒ Code of Conduct Self-Reporting Table

6 https://help.altmetric.com/support/solutions/articles/6000240580-merging-preprints-and-final-published-versions

7 https://www.altmetric.com/about-our-data/how-it-works-2/

## References

Alperin, J. P., Babini, D., & Fischman, G. (Eds.). (2014). Open access indicators and scholarly communications in Latin America. CLACSO. http://eprints.rclis.org/25122/

Alperin, J. P., Fleerackers, & Shores. (2023). Data for: Unreviewed science in the news (DRAFT VERSION) [Data set]. Harvard Dataverse. https://doi.org/10.7910/DVN/ZHQUFD

Alperin, J. P., Shores, K., & Fleerackers, A. (2023). ScholCommLab/preprints-over-time: Initial Release (1.0). Zenodo. https://doi.org/10.5281/ZENODO.8125008

Besançon, L., Peiffer-Smadja, N., Segalas, C., Jiang, H., Masuzzo, P., Smout, C., Billy, E., Deforet, M., & Leyrat, C. (2021). Open science saves lives: Lessons from the COVID-19 pandemic. BMC Medical Research Methodology, 21(1), 117. https://doi.org/10.1186/s12874-021-01304-y

Broer, I. (2020). Rapid reaction: Ethnographic insights into the Science Media Center and its response to the COVID-19 outbreak. Journal of Science Communication, 19(5), A08. https://doi.org/10.22323/2.19050208

Broer, I., & Pröschel, L. (2022). Knowledge broker, trust broker, value broker: The roles of the Science Media Center during the COVID-19 pandemic. Studies in Communication Sciences, 22(1), Article 1. https://doi.org/10.24434/j.scoms.2022.01.3070

Burke, L. (2021, January 20). Pivotal Year for Preprints. Inside Higher *Ed*. https://www.insidehighered.com/news/2021/01/20/pandemic-brought-new-attention-preprints

Chamberlain, S. (2020). Habanero [source code] (0.7.4). Github. https://github.com/sckott/habanero/releases/tag/v0.7.4

Chen, Q., Allot, A., & Lu, Z. (2020). Keep up with the latest coronavirus research. Nature, 579(7798), 193–193. https://doi.org/10.1038/d41586-020-00694-1

Deuze, M., & Witschge, T. (2018). Beyond journalism: Theorizing the transformation of journalism. Journalism: Theory, Practice & Criticism, 19(2), 165–181. https://doi.org/10.1177/1464884916688550

Else, H. (2020). How a torrent of COVID science changed research publishing—In seven charts. Nature, 588(7839), 553–553. https://doi.org/10.1038/d41586-020-03564-y

Flanagin, A., Fontanarosa, P. B., & Bauchner, H. (2020). Preprints Involving Medical Research—Do the Benefits Outweigh the Challenges? JAMA, 324(18), 1840. https://doi.org/10.1001/jama.2020.20674

Fleerackers, A., Chtena, N., Pinfield, S., Alperin, J. P., Barata, G., Oliveira, M., & Peters, I. (2023). Making science public: A review of journalists’ use of Open Science research [version 1; peer review: 1 approved]. In F1000Research (Vol. 12, Issue 512). https://doi.org/10.12688/f1000research.133710.1

Fleerackers, A., & Fagan, K. (2022). Analyzing the nature of media outlets (Version 1). OSF. https://doi.org/10.17605/OSF.IO/NW87U

Fleerackers, A., Moorhead, L. L., Maggio, L. A., Fagan, K., & Alperin, J. P. (2022). Science in motion: A qualitative analysis of journalists’ use and perception of preprints. PLOS ONE, 17(11), e0277769. https://doi.org/10.1101/2022.02.03.479041

Fleerackers, A., Nehring, L., Maggio, L. A., Enkhbayar, A., Moorhead, L., & Alperin, J. P. (2022). Identifying science in the news: An assessment of the precision and recall of Altmetric.com news mention data. Scientometrics, 127(11), 6109–6123. https://doi.org/10.1007/s11192-022-04510-7

Fleerackers, A., Riedlinger, M., Moorhead, L., Ahmed, R., & Alperin, J. P. (2021). Communicating scientific uncertainty in an age of COVID-19: An investigation into the use of preprints by digital media outlets. Health Communication, 37(6), 726–738. https://doi.org/10.1080/10410236.2020.1864892

Fraser, N., Brierley, L., Dey, G., Polka, J. K., Pálfy, M., Nanni, F., & Coates, J. A. (2021). The evolving role of preprints in the dissemination of COVID-19 research and their impact on the science communication landscape. PLOS Biology, 19(4), e3000959. https://doi.org/10.1371/journal.pbio.3000959

Fraser, N., Momeni, F., Mayr, P., & Peters, I. (2020). The relationship between bioRxiv preprints, citations and altmetrics. Quantitative Science Studies, 1(2), 618–638. https://doi.org/10.1162/qss_a_00043

Froke, P., Bratton, A. J., McMillan, J., Sarkar, P., Schwartz, J., Vadarevu, R., & Associated Press. (2020). Health, science and environment reporting. In The Associated Press stylebook 2020-2022 (55th ed.).

Funk, K. (2023, February 8). NIH preprint pilot expands to include preprints across NIH-funded research. NLM Musings from the Mezzanine. https://nlmdirector.nlm.nih.gov/2023/02/08/nih-preprint-pilot-expands-to-include-preprints-across-nih-funded-research/

Haelle, T. (2020, April 17). Beware the preprint in covering coronavirus research. Association of Health Care Journalists. https://healthjournalism.org/blog/2020/04/beware-the-preprint-in-covering-coronavirus-research/

Harrison, S., Macmillan, A., & Rudd, C. (2020). Framing climate change and health: New Zealand’s online news media. Health Promotion International, 35(6), 1320–1330. https://doi.org/10.1093/heapro/daz130

Haustein, S., Costas, R., & Larivière, V. (2015). Characterizing social media metrics of scholarly papers: The effect of document properties and collaboration patterns. PLOS ONE, 10(3), e0120495. https://doi.org/10.1371/journal.pone.0120495

Hermida, A. (2019). The existential predicament when journalism moves beyond journalism. Journalism, 20(1), 177–180. https://doi.org/10.1177/1464884918807367

Hermida, A., & Young, M. L. (2019). From peripheral to integral? A digital-born journalism not for profit in a time of crises. Media and Communication, 7(4), Article 4.

Horbach, S. P. J. M. (2020). Pandemic publishing: Medical journals strongly speed up their publication process for COVID-19. Quantitative Science Studies, 1(3), 1056–1067. https://doi.org/10.1162/qss_a_00076

Hurley, R. J., & Tewksbury, D. (2012). News aggregation and content differences in online cancer news. Journal of Broadcasting & Electronic Media, 56(1), 132–149. https://doi.org/10.1080/08838151.2011.648681

Jung, Y. E. (Grace), Sun, Y., & Schluger, N. W. (2021). Effect and reach of medical articles posted on preprint servers during the COVID-19 pandemic. JAMA Internal Medicine, 181(3), 395. https://doi.org/10.1001/jamainternmed.2020.6629

Kaiser, J. (2019, June 5). Medical preprint server debuts. Science Magazine. https://www.science.org/content/article/medical-preprint-server-debuts

Kousha, K., & Thelwall, M. (2020). COVID-19 publications: Database coverage, citations, readers, tweets, news, Facebook walls, Reddit posts. Quantitative Science Studies, 1(3), 1068–1091. https://doi.org/10.1162/qss_a_00066

Krippendorff, K. (2004). Content analysis: An introduction to its methodology (2nd ed). Sage.

Kwon, D. (2021, September 10). A Surge in Pandemic Research Shines a Spotlight on Preprints. The Scientist Magazine. https://www.the-scientist.com/news-opinion/a-surge-in-pandemic-research-shines-a-spotlight-on-preprints-69170

Maggio, L., Alperin, J. P., Moorhead, L., & Willinsky, J. (2017, March 21). Can your doctor see the cancer research reported in the news? Can you? Medium. https://medium.com/@lauren.maggio01/can-your-doctor-see-the-cancer-research-reported-in-the-news-can-you-beb9270c301f

Makri, A. (2021). What do journalists say about covering science during the COVID-19 pandemic? Nature Medicine, 27(1), 17–20. https://doi.org/10.1038/s41591-020-01207-3

Maslove, D. M. (2018). Medical preprints—A debate worth having. JAMA, 319(5), 443. https://doi.org/10.1001/jama.2017.17566

Massarani, L., & Neves, L. F. F. (2021). Reporting COVID-19 preprints: Fast science in newspapers in the United States, the United Kingdom and Brazil. Cien Saude Colet [Periódico Na Internet]. http://www.cienciaesaudecoletiva.com.br/artigos/reporting-covid19-preprints-fast-science-in-newspapers-in-the-united-states-the-united-kingdom-and-brazil/18235?id=18235

Massarani, L., Neves, L. F. F., Entradas, M., Lougheed, T., & Bauer, M. W. (2021). Perceptions of the impact of the COVID-19 pandemic on the work of science journalists: Global perspectives. Journal of Science Communication, 20(07), A06. https://doi.org/10.22323/2.20070206

Mongeon, P., & Paul-Hus, A. (2016). The journal coverage of Web of Science and Scopus: A comparative analysis. Scientometrics, 106(1), 213–228. https://doi.org/10.1007/s11192-015-1765-5

Ortega, J. L. (2020a). Altmetrics data providers: A meta-analysis review of the coverage of metrics and publication. El Profesional de La Información, 29(1). https://doi.org/10.3145/epi.2020.ene.07

Ortega, J. L. (2020b). Blogs and news sources coverage in altmetrics data providers: A comparative analysis by country, language, and subject. Scientometrics, 122(1), 555–572. https://doi.org/10.1007/s11192-019-03299-2

Penfold, N. C., & Polka, J. K. (2020). Technical and social issues influencing the adoption of preprints in the life sciences. PLOS Genetics, 16(4), e1008565. https://doi.org/10.1371/journal.pgen.1008565

Puebla, I., Polka, J., & Rieger, O. (2021). Preprints: Their Evolving Role in Science Communication [Preprint]. MetaArXiv. https://doi.org/10.31222/osf.io/ezfsk

Saari, M.-A., Gibson, C., & Osler, A. (1998). Endangered species: Science writers in the Canadian daily press. Public Understanding of Science, 7(1), 61–81. https://doi.org/10.1177/096366259800700105

Schäfer, M. S. (2017). The Oxford handbook of the science of science communication. In K. H. Jamieson, D. M. Kahan, & D. A. Scheufele (Eds.), How Changing Media Structures Are Affecting Science News Coverage (Vol. 1). Oxford University Press. https://doi.org/10.1093/oxfordhb/9780190497620.013.5

Schapals, A. K. (2022). Peripheral actors in journalism: Deviating from the norm? Routledge (Taylor & Francis). https://doi.org/10.4324/9781003144663

Schultz, T. (2023). A survey of U.S. science journalists’ knowledge and opinions of open access research. International Journal of Communication, 17, Article 0.

Schwab, L. (2021). arxiv: Python wrapper for the arXiv API: http://arxiv.org/help/api/ (1.4.2) [Python; OS Independent]. 0.7.4. https://github.com/lukasschwab/arxiv.py

Simons, A., & Schniedermann, A. (2023). Preprints in the German news media before and during the COVID pandemic. A comparative mixed-method analysis. In I. Broer, S. Lemke, A. Mazarakis, I. Peters, & C. Zinke-Wehlmann (Eds.), The Science-Media Interface: On the Relation Between Internal and External Science Communication (pp. 53–77). De Gruyter Saur. https://www.degruyter.com/document/isbn/9783110776546/html?lang=en

StataCorp. (2021). Stata Statistical Software (Version 17). StrataCorp LLC

Stocking, G. (2019). Trends and facts on online news: State of the news media. Pew Research Center. https://www.journalism.org/fact-sheet/digital-news/

The Pandas Development Team. (2023). Pandas (1.5.3) [Python]. https://doi.org/10.5281/zenodo.7549438

van Schalkwyk, F., & Dudek, J. (2022). Reporting preprints in the media during the Covid-19 pandemic: Supplemental material. Public Understanding of Science, 2.

Vergoulis, T., Kanellos, I., Chatzopoulos, S., Pla Karidi, D., & Dalamagas, T. (2021). BIP4COVID19: Releasing impact measures for articles relevant to COVID-19. Quantitative Science Studies, 2(4), 1447–1465. https://doi.org/10.1162/qss_a_00169

Waltman, L., Pinfield, S., Rzayeva, N., Henriques, S. O., Fang, Z., Brumberg, J., Greaves, S., Hurst, P., Collings, A., Heinrichs, A., Lindsay, N., MacCallum, C., Morgan, D., Sansone, S.-A., & Swaminathan, S. (2021). Scholarly communication in times of crisis [Report]. Research on Research Institute. https://apo.org.au/node/315479

WHO. (2020, March 19). Infection prevention and control during health care when novel coronavirus (nCoV) infection is suspected. WHO. https://www.who.int/publications-detail-redirect/10665-331495

Wihbey, J. (2017). Journalists’ use of knowledge in an online world: Examining reporting habits, sourcing practices and institutional norms. Journalism Practice, 11(10), 1267–1282. https://doi.org/10.1080/17512786.2016.1249004

Witschge, T., Anderson, C., Domingo, D., & Hermida, A. (2019). Dealing with the mess (we made): Unraveling hybridity, normativity, and complexity in journalism studies. Journalism, 20(5), 651–659. https://doi.org/10.1177/1464884918760669

